# Development of a genetically encoded supersulfide-dependent translocation reporter

**DOI:** 10.64898/2026.06.17.733027

**Authors:** Shouhei Misaki, Tatsuya Kandaka, Takashi Tanida, Shingo Kasamatsu, Tomoya Ito, Hideshi Ihara, Yasu-Taka Azuma, Motohiro Nishida, Kazuhiro Nishiyama

## Abstract

Supersulfides are emerging sulfur-containing signaling molecules involved in redox regulation, mitochondrial function, and protein S-sulfhydration. However, their dynamic behavior in living mammalian systems remains poorly understood because existing analytical methods require destructive sample preparation or lack sufficient intracellular applicability. Here, we developed a genetically encoded supersulfide-dependent translocation reporter (SuTR) for mammalian cells and in vivo imaging. Although the previously reported probe psGFP failed to respond to supersulfides in mammalian cells, fusion of psGFP with the sulfide-responsive transcriptional repressor (SqrR) generated SuTR, a novel reporter that exhibited supersulfide-dependent translocation from the nucleus to the cytoplasm. Na₂S₂ and Na₂S₃ induced dose-dependent cytosolic translocation of SuTR, whereas Na₂S showed no effect. Fluorescence recovery after photobleaching (FRAP) analysis revealed accelerated fluorescence recovery shortly after supersulfide stimulation, and overexpression of the endogenous supersulfide-producing enzyme Cysteinyl-tRNA Synthetase 2 (CARS2) similarly altered reporter dynamics. Mutational analyses demonstrated that reporter responsiveness depends on the DNA-binding activity of SqrR. Furthermore, SuTR successfully detected supersulfide induction in mouse liver in vivo following Na₂S₃ administration. These findings establish SuTR as a genetically encoded reporter for monitoring supersulfide dynamics in mammalian cells and tissues.

**Highlights:**

- We developed SuTR, a genetically encoded supersulfide-dependent translocation reporter.
- Supersulfides induce nuclear-to-cytoplasmic translocation of SuTR
- FRAP enables rapid detection of endogenous and exogenous supersulfide responses
- SuTR activity depends on the DNA-binding function of SqrR
- SuTR enables visualization of supersulfide dynamics in mouse liver in vivo

## 1. Introduction

Supersulfides, collectively referred to as reactive sulfur species (RSS), have emerged as important regulators in redox biology. Characterized by a sulfane sulfur chain structure (RSSnR, where R represents hydrogen or an alkyl group and n > 1), these molecules possess markedly greater redox reactivity than conventional sulfur-containing metabolites, including cysteine, glutathione, and hydrogen sulfide (H_2_S)[1,2]. Growing evidence suggests that these sulfur-containing molecules are involved in a wide range of cellular functions, including the modulation of Ca²⁺ signaling, regulation of tumor suppressor phosphatases, protein S-sulfhydration and preservation of intracellular redox homeostasis [3–10]. In addition, supersulfides have been implicated in the control of mitochondrial function, including mitochondrial biogenesis and energy metabolism [11].

Despite increasing recognition of the biological importance of supersulfides [12–14], their spatiotemporal dynamics in living cells and tissues remain poorly understood. A major obstacle is the lack of tools capable of monitoring supersulfides in real time within intact biological systems. Consequently, fundamental questions regarding where, when, and under what physiological or pathological conditions supersulfides are generated remain largely unanswered.

Mass spectrometry-based supersulfideomics has enabled sensitive detection and quantification of endogenous supersulfides [15–17]. However, because these analyses require cell disruption, they are not suitable for real-time imaging or spatial analysis in intact cells [18]. As a result, dynamic changes in supersulfide signaling cannot be directly visualized. Fluorescent chemical probes, including SSP4, Ssip-1 and QS10, have been established for the detection of supersulfides[19–21]. Despite their usefulness in cultured cell systems, chemical probes generally require exogenous loading and may suffer from limited tissue penetration, making long-term monitoring in living animals challenging.

Genetically encoded fluorescent reporters represent an attractive strategy for overcoming these limitations because they can be continuously expressed in living cells and animals, enabling long-term monitoring of intracellular signaling events. However, currently available genetically encoded supersulfide reporters have been primarily characterized in microorganisms, and their applicability to mammalian cells and tissues remains uncertain[22,23]. We therefore sought to develop a genetically encoded supersulfide reporter suitable for mammalian cells and in vivo imaging. Such a tool would facilitate direct visualization of supersulfide dynamics in living tissues and may provide a platform for uncovering previously inaccessible aspects of supersulfide biology in health and disease.

In the present study, we developed and evaluated a genetically encoded supersulfide-dependent translocation reporter (SuTR) optimized for use in mammalian cells and in vivo imaging applications.

## 2. Materials and Methods

### 2.1. Cell culture

Human embryonic kidney cells (HEK293) and HeLa cells were maintained in Dulbecco’s modified Eagle’s medium (DMEM) containing 10% fetal bovine serum (FBS) and 1% penicillin–streptomycin under standard culture conditions.

### 2.2. Plasmid construction

The SuTR expression constructs were synthesized by VectorBuilder. Site-specific amino acid substitutions were introduced by PCR-based mutagenesis. The coding sequence of mouse CARS2 was amplified from mouse cDNA by PCR and cloned into the pTS2387-Tier1-PhCMV-p2a_mTagBFP2 vector using the NEBuilder HiFi DNA Assembly Cloning Kit (New England Biolabs) to generate the CARS2 expression construct. pTS2387-Tier1-PhCMV-p2a_mTagBFP2 was a gift from Martin Fussenegger (Addgene plasmid # 169524 ; http://n2t.net/addgene:169524 ; RRID:Addgene_169524) [24]. Details of the SuTR sequence and PCR primers used in this study are summarized in Supplementary Table 1 and 2.

### 2.3. Plasmid transfection

HEK293 and HeLa cells were transiently transfected with plasmid DNA using ViaFect Transfection Reagent (Promega) following the supplier’s protocol.

### 2.4. Live cell imaging

HEK293 or HeLa cells expressing SuTR were seeded onto glass coverslips prior to imaging. Time-lapse fluorescence imaging was conducted using Zeiss LSM980 confocal laser-scanning microscope or FV3000 confocal laser-scanning microscope (Olympus) and a stage-top incubation system maintained at 37℃ with 5% CO₂. GFP fluorescence was excited using a 488-nm argon laser. Na_2_S (100 μM), Na_2_S_2_ (100 μM), or Na_2_S_3_ (100 μM) were added, and cells were imaged every 15 or 30 min for 90 min in Live Cell Imaging Solution (Thermo Fisher Scientific).

To investigate the concentration dependence, Na_2_S (0-100 μM), Na_2_S_2_ (0-100 μM), or Na_2_S_3_ (0-100 μM) was added to cells, fixed with 4% PFA after 90 min, and GFP fluorescence was observed.

### 2.5. Fluorescence recovery after photobleaching (FRAP) Analysis

FRAP experiments were conducted using an FV3000 confocal microscope as previously described [25]. HeLa cells were plated onto 35-mm glass-bottom dishes at a density of 3 × 10⁴ cells/dish and transfected with the SuTR construct as described above. Following transfection, fluorescence recovery was monitored after selective photobleaching of GFP signals. Images were acquired using a 40×objective under the following settings: 512 × 512-pixel resolution, zoom factor 10, scan speed 2 μsec/pixel, and laser intensity corresponding to 100% of maximum power. GFP bleaching was performed with a 488-nm laser at full power for 200 μsec/pixel. Time-lapse images were subsequently captured every 2 sec, with the initial post-bleach frame designated as 0 sec. For each experiment, a circular region of interest (ROI: 12 pixels) was selected within randomly chosen transfected cells and subjected to photobleaching. Fluorescence recovery within a smaller measurement area was quantified at each time point using FV3000 analysis software. Recovery kinetics were evaluated from 20 independent cells, and the half-time of fluorescence recovery (T_1/2_) was calculated from the corresponding recovery curves.

### 2.6. Structure prediction using AlphaFold 3

The amino acid sequence of SqrR was obtained from a previous report [26] and submitted to the AlphaFold 3 platform[27] via the AlphaFold Server (https://alphafoldserver.com/). To predict the homodimeric structure, the SqrR sequence was entered as the sole protein component, and the copy number was set to two, while all other parameters were retained at their default settings. Structure prediction was performed using the cloud-based AlphaFold Server. For protein–DNA complex modeling, DNA sequences reported previously [26] (5′-TTGACAGGTTGCCTATTCATATTCTCATATGTGC-3′ and 5′-GCACATATGAGAATATGAATAGGCAACCTGTCAA-3′) were included through the AlphaFold 3 interface. Predicted structures were visualized using PyMOL v3.1 or by directly examining images generated by the AlphaFold Server.

### 2.7 Animals

All animal procedures were performed in compliance with institutional regulations and the National Institutes of Health Guide for the Care and Use of Laboratory Animals.

The study protocols received approval from the Animal Care and Use Committee of Osaka Metropolitan University (approval numbers: 25–146, approved December 8th, 2025; and 26–41, approved April 1, 2026). All experiments involving animals were conducted in accordance with the ARRIVE guidelines [28].

Male C57BL/6 mice aged 8–10 weeks and weighing 19–23 g were obtained from CLEA Japan (Tokyo, Japan). Animals were maintained in individually ventilated cages containing wood-chip bedding under standardized conditions, including a 12-h light/dark cycle, ambient temperature of 21–23℃, and relative humidity of 50–60%. Food and water were available ad libitum throughout the study.

### 2.8. AAV

Recombinant AAV particles were produced in HEK293T cells by transient co-transfection of the transfer plasmid together with the pAAV2/8 packaging vector and pHelper plasmid. Cell culture supernatants were collected beginning 48 h after transfection, and harvested cells were combined with the medium at 96 h post-transfection.Virus purification was carried out based on an established procedure [29]. Briefly, chloroform extraction was performed to eliminate cellular contaminants and insoluble materials, after which viral particles were precipitated using PEG8000. The precipitated fraction was resuspended in 50 mM HEPES buffer and subjected to an additional chloroform purification step. Concentration and buffer exchange into PBS were achieved using 100-kDa molecular weight cutoff centrifugal filters.

### 2.9. In vivo cell imaging

Mice were anesthetized by intraperitoneal administration of a medetomidine–midazolam–butorphanol mixture (0.3, 4, and 5 mg/kg, respectively). Under anesthesia, AAV vectors were administered via the retro-orbital sinus at a dose of 2 × 10¹¹ vector genomes per mouse.

Three weeks after AAV administration, mice received intraperitoneal injections of Na₂S₃ (20 mg/kg) once daily for 5 consecutive days. In a separate experiment, oxidative stress was induced by a single intraperitoneal injection of thioacetamide (TAA, 100 mg/kg). Mice were euthanized 6 h after TAA administration.

At the experimental endpoint, mice were deeply anesthetized by intraperitoneal administration of a medetomidine–midazolam–butorphanol mixture (0.3, 4, and 5 mg/kg, respectively) and euthanized by exsanguination via the inferior vena cava. Liver tissues were collected and fixed with 4% paraformaldehyde. Following embedding in optimal cutting temperature (OCT) compound (Sakura Finetek), samples were snap-frozen and sectioned at a thickness of 10 μm using a cryostat. GFP fluorescence was visualized using a confocal laser-scanning microscope (instrument model).

Fluorescence intensities in nuclear and cytoplasmic compartments were quantified using ImageJ software. Cells were randomly selected for analysis, and 10–15 cells per animal were evaluated. Nuclear-to-cytoplasmic GFP fluorescence ratios were averaged for each mouse and used for statistical analysis. Both vehicle-treated and Na₂S₃-treated groups consisted of five animals.

### 2.10. Measurement of supersulfide levels by LC–ESI–MS/MS following TME-IAM labeling

Quantitative analysis of supersulfide metabolites was conducted using TME-IAM derivatization as previously described [30], with minor modifications. In brief, plasma samples were incubated with 1 mM TME-IAM at 30℃ for 30 min. The reaction mixtures were subsequently diluted fourfold with 0.1% formic acid (Nacalai Tesque, Inc.) and supplemented with stable isotope-labeled TME-AM adduct standards at known concentrations prepared in 0.1% FA. Quantification of TME-IAM-derivatized metabolites was subsequently performed by LC–ESI–MS/MS using an LCMS-8060 system following a previously established protocol [30].

### 2.11. Materials

Na₂S_2_, Na₂S₃, Na₂S_4_ were obtained from Dojindo (Kumamoto, Japan). Na_2_S was purchased from Tokyo Chemical Industry (Tokyo, Japan). H_2_O_2_ and TAA were purchased from Nacalai Tesque (Kyoto, Japan). Leptomycin B was purchased from Cayman Chemical.

### 2.12. Statistics

Data are presented as mean ± SEM from a minimum of three biologically independent experiments. Differences were considered statistically significant at P < 0.05. For comparisons between two groups, unpaired Student’s t-tests were applied.

Multiple-group analyses were conducted using one-way ANOVA followed by Tukey’s post hoc test or two-way ANOVA with Sidak’s post hoc test. Statistical calculations were carried out using GraphPad Prism version 9.0 (GraphPad Software).

## 3. Results

### 3.1. Development of translocation reporter for detection of supersulfide

We first attempted to detect supersulfide in mammalian cells using psGFP[22]. However, the fluorescence of psGFP did not change even when the supersulfide donor Na_2_S_3_ (200 μM) was added (Supplementary Figure 1). We therefore sought to enhance the responsiveness of psGFP in mammalian cells. The redox probe roGFP can be fused to redox-sensitive Grx1 or Orp1 to increase its sensitivity and specificity [31,32]. Therefore, we fused the supersulfide-sensitive protein, Sulfide-responsive transcriptional repressor (SqrR) to psGFP.

Unexpectedly, although fusion of SqrR did not enhance the fluorescence responsiveness of psGFP, the resulting construct exhibited supersulfide-dependent translocation from the nucleus to the cytoplasm. SqrR-psGFP expressed in mammalian cells was predominantly localized in the nucleus, and Na_2_S_3_ (200 μM) treatment induced its redistribution to the cytoplasm without altering fluorescence intensity (Figure 1A, B and Supplementary Figure 2). We therefore designated this construct a supersulfide-dependent translocation reporter (SuTR). The administration of supersulfide donors Na_2_S_3_ and Na_2_S_2_ dose-dependently shifted the localization of SuTR from the nucleus to the cytoplasm, whereas the administration of H_2_O_2_ or Na_2_S did not alter the localization of SuTR, which remained exclusively in the nucleus (Figure 1C-F). These findings indicate that SuTR functions as a genetically encoded reporter for supersulfide signaling based on nucleocytoplasmic translocation.

**Figure 1.**
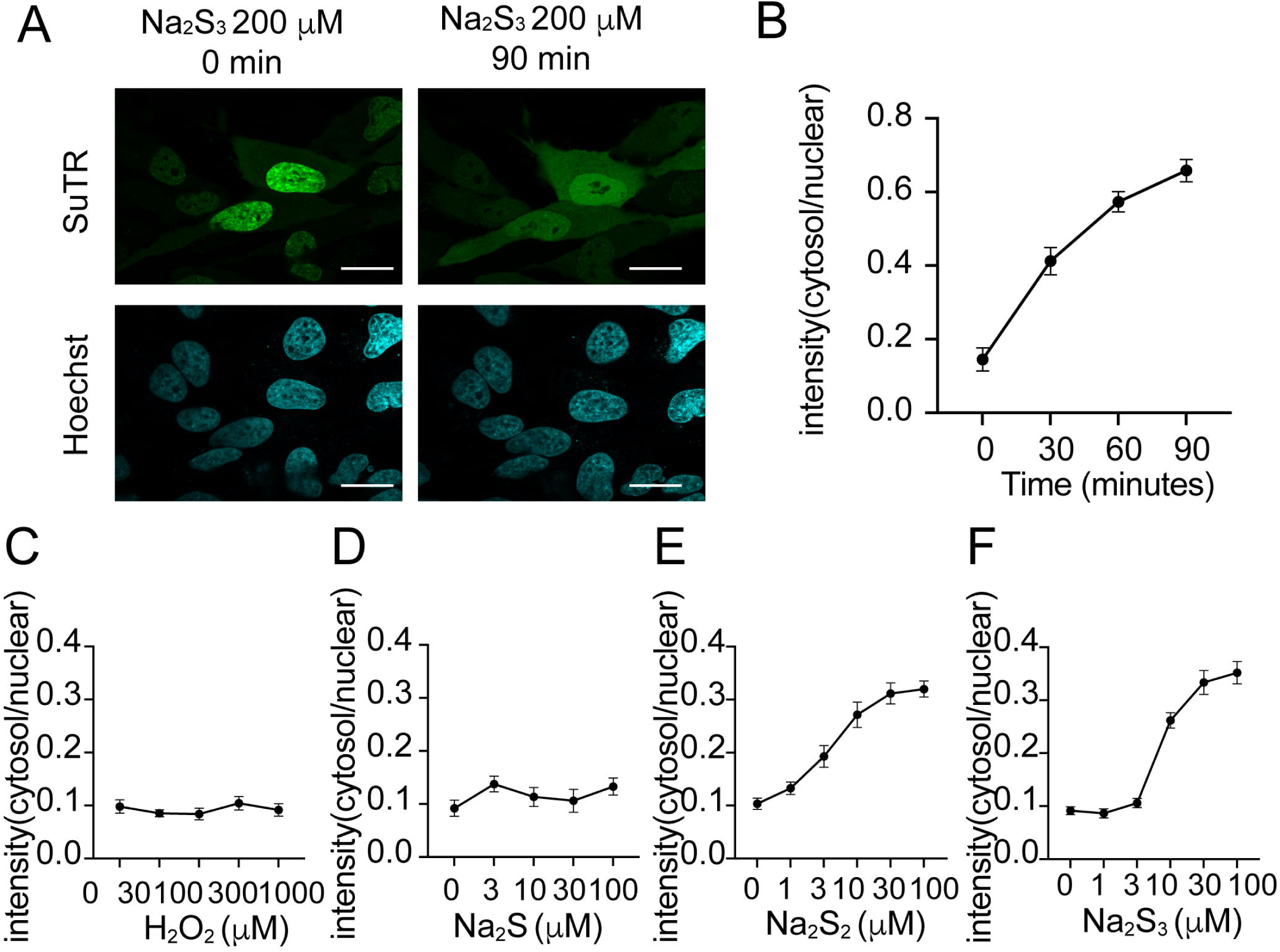
Development of translocation reporter for detection of supersulfide. (A) HeLa cells expressing SuTR were stimulated with Na_2_S_3_ (200 μM) and imaged at indicated time points. Scale bar, 20 μm (B) HeLa cells were stimulated with Na_2_S_3_ (200 μM), imaged, and quantified. Data are represented as the mean ± SEM from 13 cells. HeLa cells were stimulated with H_2_O_2_ (0-1000 μM) (C) Na_2_S (0-100 μM) (D), Na_2_S_2_ (0-100 μM) (E), Na_2_S_3_ (0-100 μM) (F), imaged, and quantified. Data are shown as the mean ± SEM from 20-87 cells. A higher concentration (200 μM) was used in the initial time-course experiment to facilitate visualization of reporter translocation, whereas subsequent concentration–response analyses were performed using 0–100 μM.

### 3.2. FRAP analysis of SuTR

We found that it takes about 90 min for SuTR to translocate from the nucleus to the cytoplasm. Therefore, we performed FRAP analysis to characterize early supersulfide-dependent changes in SuTR dynamics. Administration of Na_2_S_2_ (10 μM and 100 μM) significantly reduced the time required for fluorescence recovery (T_1/2_) (Figure 2A-C). Overexpression of Cysteinyl-tRNA Synthetase 2 (CARS2), a supersulfide-producing enzyme [33], shortened the fluorescence recovery time, accompanied by a decrease in the mobile fraction (Figure 3A, B). These findings demonstrate that both exogenous and endogenous supersulfides can be rapidly detected using SuTR in combination with the FRAP assay.

**Figure 2.**
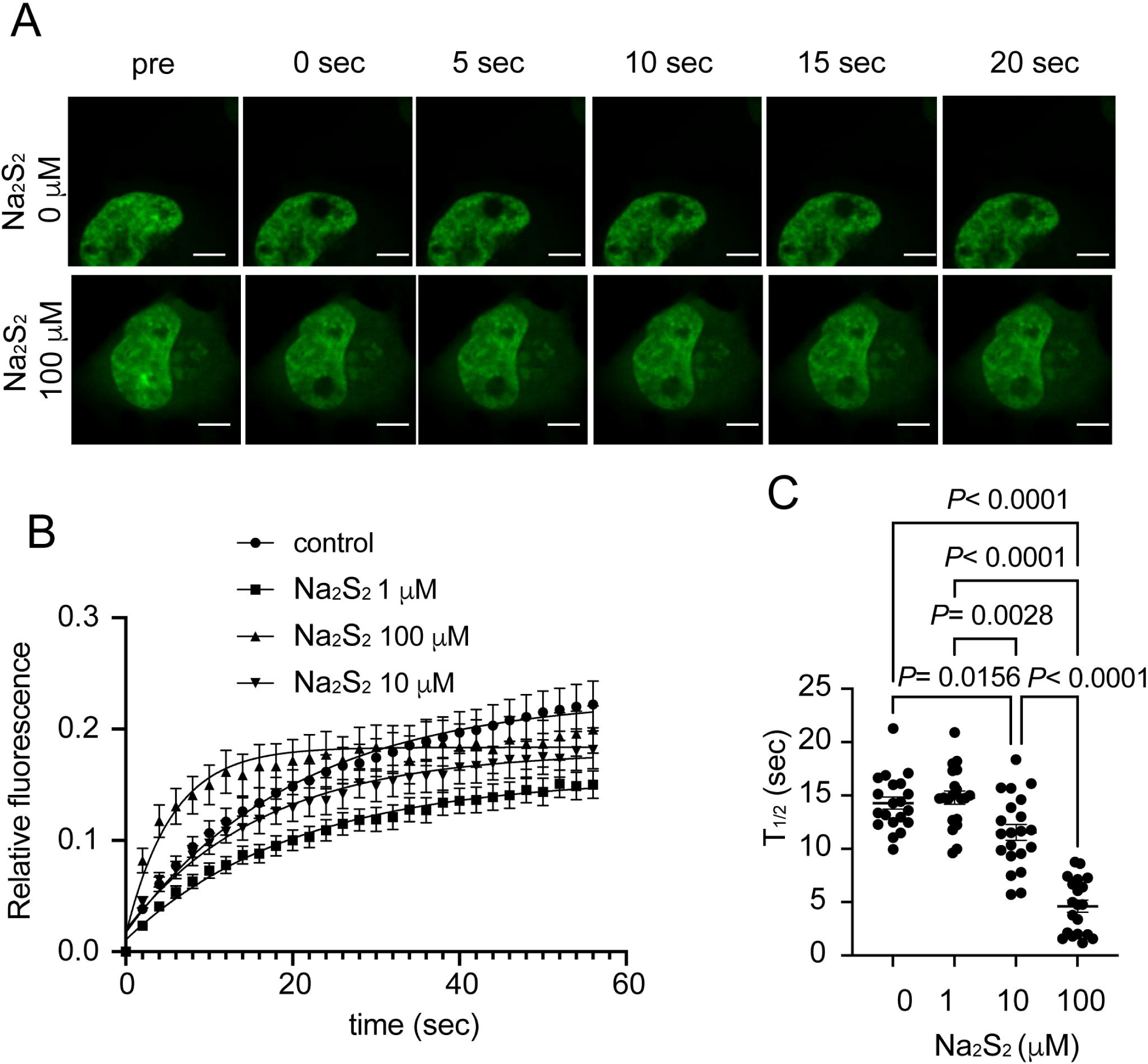
FRAP analysis of SuTR in HeLa cells. (A) HeLa cells expressing SuTR were stimulated with Na_2_S_2_ (100 μM) and imaged at indicated time points. Scale bar, 5 μm (B) GFP recovery curves of FRAP for cells expressing SuTR treated with Na_2_S_2_ (100 μM). The time of initial fluorescence intensity after bleaching was set to 0 sec. (C) Quantification of FRAP analyses. Data are shown as the mean ± SEM from 20-21 cells. Statistical significance was determined using one-way ANOVA followed by Tukey’s multiple-comparison test.

**Figure 3.**
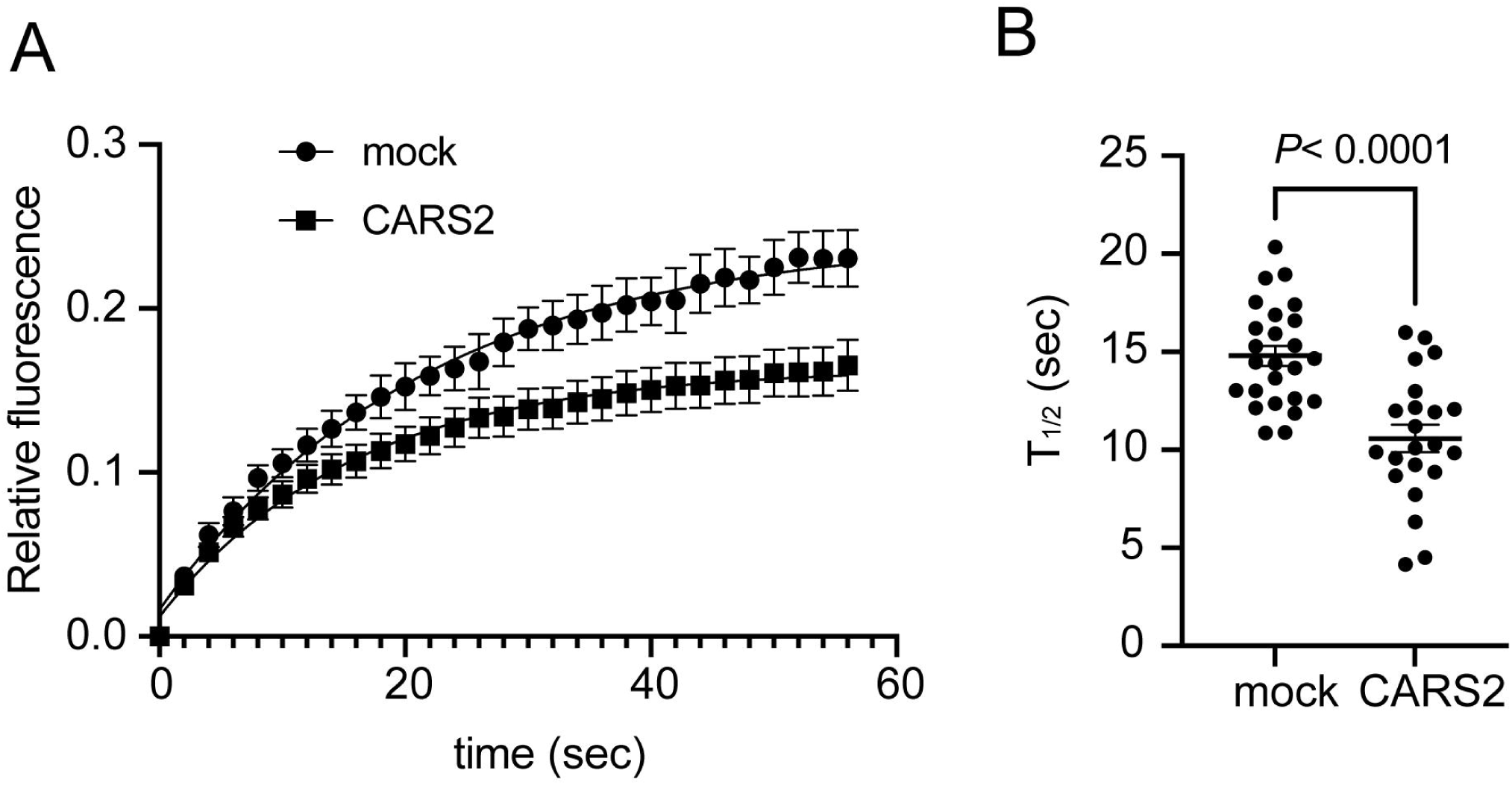
CARS2 overexpression enhances SuTR translocation dynamics. (A) Co-transfection with SuTR and CARS2 or mock was carried out in HeLa cells. FRAP analyses of GFP were performed. GFP recovery curves of FRAP for cells co-expressing SuTR and CARS2 or mock. (B) Quantification of FRAP analyses. Data are shown as the mean ± SEM from 22-25 cells. Significance was determined using unpaired *t*-test.

### 3.3. The DNA-binding domain of SqrR affects the localization of SuTR

SqrR is a repressor that binds to DNA when the concentration of supersulfide is low, and its binding to DNA weakens as the concentration of supersulfide increases [26,34]. The C9S mutant, in which the ninth cysteine residue is replaced with serine, is known to have increased DNA binding [26,34]. Furthermore, α-helix 4 is predicted to be the DNA binding site[34] (Figure 4A, B). Therefore, we created the C9S mutant and mutants with amino acid substitutions in α-helix 4 (L68A, R70A, R72A) and observed their intracellular localization. The WT was localized in the nucleus and was relocated to the cytoplasm by Na_2_S_2_ treatment, whereas the C9S mutant was localized in the nucleus, similar to the WT, but did not change its localization even with Na_2_S_2_ treatment. The α-helix 4 mutants were distributed throughout both the cytoplasm and nucleus, and their localization was not altered by Na_2_S_2_ administration (Figure 4C, D). Treatment with Leptomycin B, an inhibitor of Exportin 1 (XPO1)-mediated nuclear export, failed to suppress the supersulfide-induced translocation of SuTR (Supplementary Figure 3A, B). These findings suggest that the nuclear localization of SqrR in mammalian cells depends on its DNA-binding activity and that supersulfide-induced weakening of DNA binding results in the loss of nuclear retention, leading to its redistribution to the cytoplasm.

**Figure 4.**
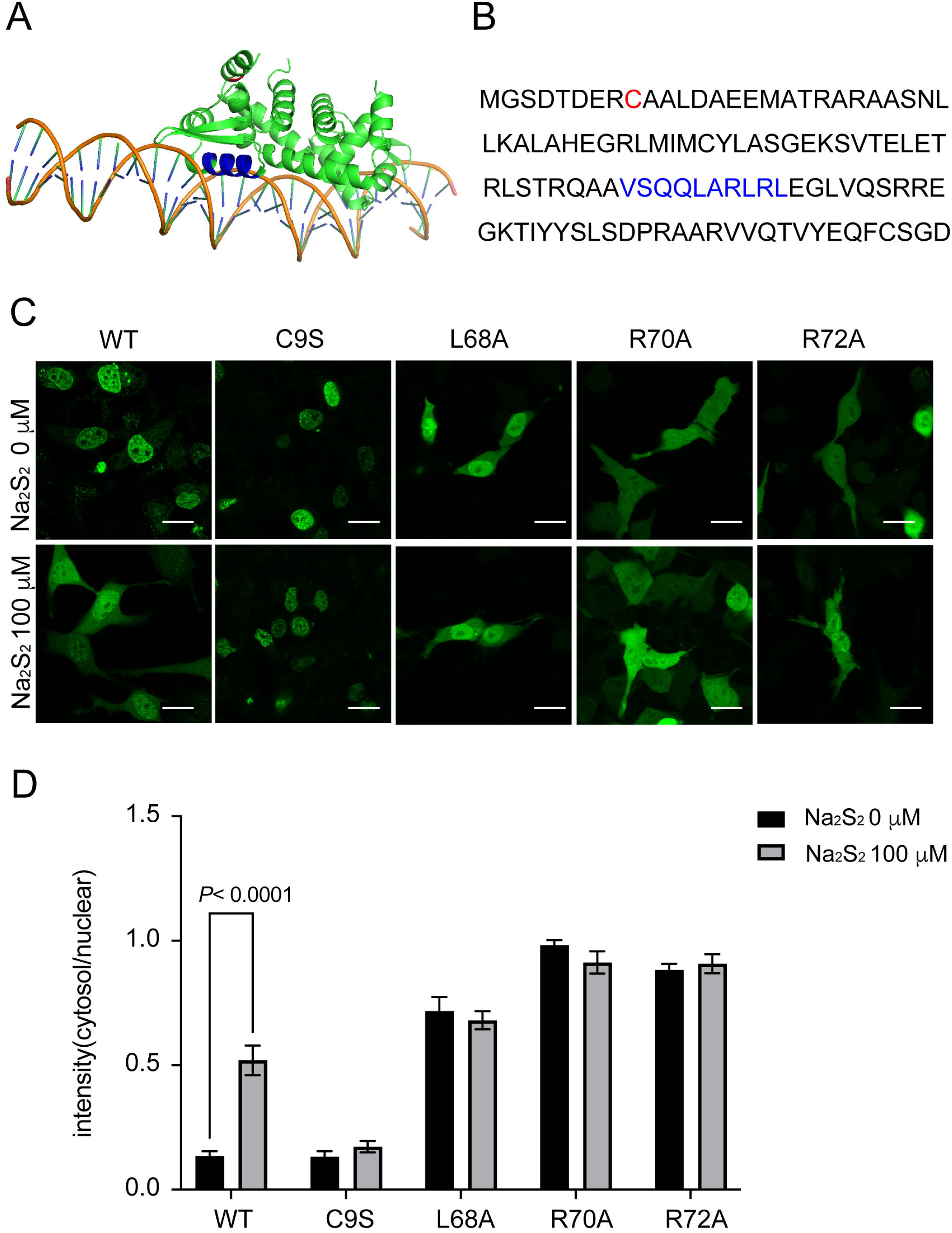
The DNA-binding domain of SqrR affects the localization of this reporter. (A) Predictive structural model of SqrR-DNA complex using AlphaFold 3. The ninth cysteine residue is highlighted in red, whereas α-helix 4 is shown in blue. (B) Sequence alignment of SqrR. (C) HeLa cells expressing SuTR mutants were stimulated with Na_2_S_2_ (100 μM). Scale bar, 20 μm (D) Quantification of the nuclear and cytoplasmic fluorescence intensity ratio of SuTR. Data are shown as the mean ± SEM from 20 cells. Significance was determined using two-way ANOVA with Sidak’s comparison test.

### 3.4. Verification of SuTR reactivity in mouse liver

To investigate the in vivo response of SuTR to supersulfide induction, mice expressing SuTR were treated with Na₂S₃. Repeated administration of Na₂S₃ (20 mg/kg) resulted in clear translocation of the reporter from the nucleus to the cytoplasm, consistent with the in vitro observations described above. Quantitative analysis revealed a significant increase in the cytoplasmic-to-nuclear GFP fluorescence intensity ratio in Na₂S₃-treated animals compared with vehicle controls (Figure 5A, B). Na_2_S_3_ administration increased plasma concentrations of CysSH, CysSSH, GSH, and GSSH (Figure 5C). On the other hand, SuTR did not respond to TAA-induced liver injury (Figure 5A, B). These findings demonstrate that SuTR can detect supersulfide induction in mouse liver in vivo.

**Figure 5.**
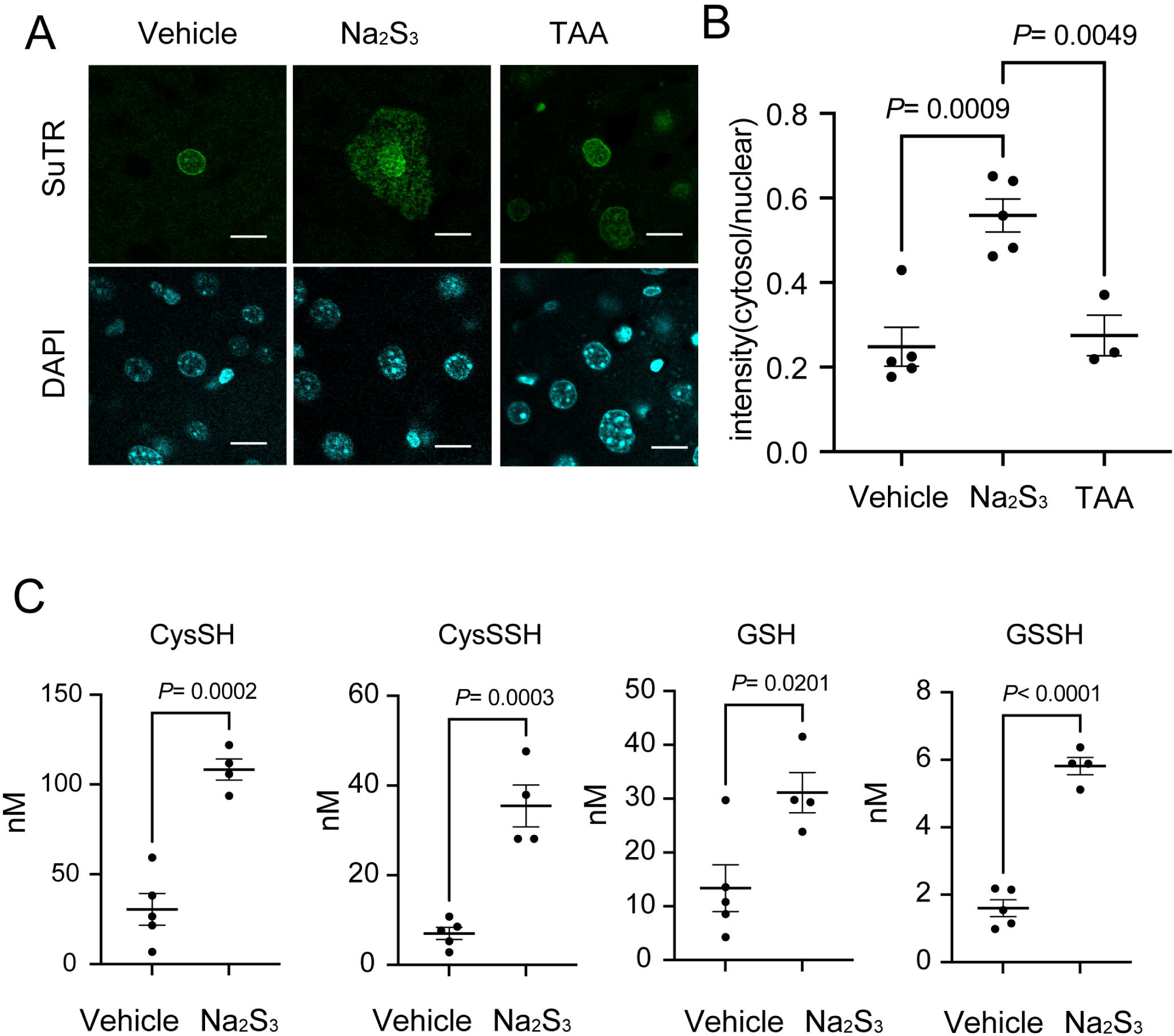
In vivo validation of SuTR in mouse liver. After expressing SuTR using AAV vectors, mice were treated with saline, Na_2_S_3_ (20 mg/kg, i.p. for 5 days) or TAA. GFP fluorescence images of fixed liver tissue. (A) representative images. Scale bar, 20 μm (B) Quantification of the ratio of fluorescence intensity between the nucleus and cytoplasm. Data are shown as the mean ± SEM (N = 5 mice for Vehicle, N = 5 mice for Na_2_S_3_, N = 3 for TAA). (C) Levels of CysSH, GSH, CysSSH, and GSSH in the plasma of mice . Data are shown as the mean ± SEM (N = 5 mice for Vehicle, N = 4 mice for Na_2_S_3_). Significance was determined using one-way ANOVA followed by Tukey’s multiple-comparison test (B) or unpaired *t*-test (C).

## 4. Discussion

In this study, we developed SuTR, a genetically encoded reporter that enables visualization of supersulfide dynamics in mammalian cells and tissues. By overcoming key limitations of existing analytical approaches, this reporter provides a platform for investigating the spatiotemporal regulation of supersulfide signaling in vivo.

Previously, roGFP and psGFP have been reported as redox-responsive protein-based probes [22,35]. Because roGFP alone has low sensitivity and specificity, its fusion with Grx1 or Orp1 enhances these functions [31,32]. Supersulfide-responsive protein probes, such as psGFP and psRFP, have been developed based on roGFP [22,23].

However, these probes have only been used in bacteria and yeast, not mammalian cells. Therefore, we expressed psGFP in mammalian cells and administered supersulfides, but they did not respond. It has also been reported that psRFP does not work well in mammalian cells. This may be due to the stronger buffering effect of redox in mammalian cells compared to bacteria and yeast [36,37]. To enhance the sensitivity and specificity of psGFP, we fused it to SqrR, a transcription factor responsive to supersulfides. SqrR is a transcription factor identified as a master regulator of sulfide metabolism in the purple bacterium Rhodobacter capsulatus [26]. Two cysteine residues (Cys41 and Cys107) conserved among SqrR homologs form an intramolecular tetrasulfide bond in a supersulfide-dependent manner, reducing the DNA-binding affinity of SqrR. Aiming to enhance the sensitivity or specificity of psGFP, we constructed SuTR and expressed it in mammalian cells. Unexpectedly, fusion of SqrR to psGFP did not enhance fluorescence responsiveness but instead generated a supersulfide-dependent nuclear-to-cytoplasmic translocation response. No mammalian homolog of SqrR has been identified to date. However, the nuclear localization of SuTR and the mutational analyses suggest that SqrR interacts with mammalian DNA and that supersulfides weaken this interaction, similar to the mechanism reported in purple bacteria. The SuTR responded to 3-100 μM Na_2_S_2_ or Na_2_S_3_, demonstrating responsiveness within a concentration range comparable to that reported for QS10. However, the translocation of SuTR from the nucleus to the cytoplasm takes approximately 60 to 90 min. To investigate early supersulfide-dependent changes in SuTR behavior, we performed FRAP analysis. The fluorescence recovery of SuTR after photobleaching was accelerated 10 min after Na_2_S_2_ administration. FRAP analysis revealed a rapid increase in SuTR mobility following supersulfide stimulation. Because fluorescence recovery is influenced by both molecular diffusion and binding interactions, these observations do not directly demonstrate dissociation from DNA. However, together with the mutational analyses showing a requirement for the DNA-binding region of SqrR, the FRAP results are consistent with a model in which supersulfides reduce DNA binding and thereby increase the mobility of SuTR within the nucleus.

Furthermore, existing probes were based on fluorescence intensity changes, whereas SuTR is a translocation-based reporter. An advantage of SuTR is that reporter activity can be evaluated in both live-cell imaging and fixed specimens. Unlike intensity-based probes such as roGFP, which can be engineered to monitor redox responses within specific organelles by incorporating organelle-targeting signals, SuTR cannot be adapted for organelle-specific measurements because its sensing mechanism relies on nucleocytoplasmic translocation.

The Kinase Translocation Reporter (KTR) platform comprises reporters for multiple MAP kinase pathways, including ERK1/2, p38, and JNK. Upon pathway activation, these reporters undergo nucleocytoplasmic translocation, relocating from the nucleus to the cytoplasm in response to kinase signaling [38–41]. Many translocation reporters have been developed based on mammalian proteins. SuTR, developed in this study, is a rare translocation reporter derived from a bacterial protein. The successful use of a bacterial transcription factor as the sensing module suggests that stress-responsive proteins from microorganisms may represent a valuable resource for developing genetically encoded translocation reporters in mammalian systems.

We further demonstrated that SuTR can be applied to mouse tissues in vivo. Typically, genetically encoded probes based on fluorescence changes require a multiphoton microscope to observe deep tissue when used in tissues such as those of mice. Because SuTR is a probe that uses translocation as an indicator, it does not require expensive equipment like a multiphoton microscope and can be observed using a fluorescence microscope with widely used tissue sections. An additional advantage of the translocation-based design is that reporter activity can be quantified in fixed tissue sections, facilitating in vivo applications that are difficult to achieve with many intensity-based fluorescent sensors. Furthermore, in this study, a TAA-induced liver injury model was used as a model of oxidative stress in the liver. Although TAA itself is not hepatotoxic, it is converted to the toxic metabolite TAA-S-oxide by the liver’s cytochrome P450 enzymes [42,43]. The resulting metabolite generates a large amount of reactive oxygen species (ROS) in hepatocytes [43]. SuTR did not respond to hydrogen peroxide treatment or TAA-induced oxidative stress. These findings support the selectivity of SuTR toward supersulfides over the oxidative stress conditions examined in this study.

A limitation of SuTR is that reporter activation relies on nucleocytoplasmic translocation and is therefore slower than fluorescence intensity-based sensors. In addition, the current design is optimized for detecting increases in supersulfide levels and may not be suitable for monitoring supersulfide depletion. Future studies will be required to determine whether the reporter can detect endogenous supersulfide fluctuations under physiological and pathological conditions. Future studies using SuTR may reveal how endogenous supersulfides are dynamically regulated during physiological adaptation and disease progression in vivo.

## 5. Conclusion

In the present study, we developed and evaluated a genetically encoded supersulfide reporter optimized for use in mammalian cells and in vivo imaging applications.

Currently, quantitative methods using mass spectrometry, imaging using fluorescent chemical probes, and imaging methods using Raman spectroscopy have been established [16,21,44,45]. However, none of these methods fully capture the dynamics of supersulfides in living organisms (tissues). Our findings establish SuTR as the first genetically encoded supersulfide-dependent translocation reporter applicable to mammalian cells and tissues.

## Acknowledgments

We sincerely thank Dr. Akiyuki Nishimura and Dr. Yuri Kato for helpful comments. This work was supported by JST CREST Grant Number JPMJCR2024 (20348438 to M.N), JSPS KAKENHI (24KK0275 and 22K06630 to K.N., and 22K19395 to M.N.), Grant-in-Aid for Scientific Research on Innovative Areas(A) “Sulfur biology” (24H01326 to K.N., and 21H05269 to M.N.) from the Ministry of Education, Culture, Sports, Science and Technology of Japan.

## CRediT authorship contribution statement

**Shouhei Misaki:** Writing – review & editing, Writing – original draft, Data curation, Conceptualization. **Tatsuya Kandaka:** Writing – review & editing, Data curation. **Takashi Tanida:** Writing – review & editing, Resources, Methodology. **Shingo Kasamatsu:** Writing – review & editing, Resources, Methodology. **Tomoya Ito:** Writing – review & editing, Data curation. **Hideshi Ihara:** Writing – review & editing, Resources, Methodology. **Yasu-Taka Azuma:** Writing – review & editing, Resources. **Motohiro Nishida:** Writing – review & editing, Funding acquisition. **Kazuhiro Nishiyama:** Writing – review & editing, Writing – original draft, Funding acquisition, Data curation, Conceptualization.

## Data Availability Statement

The data supporting the findings of this study are available from the corresponding author upon reasonable request.

## Declarations

During the preparation of this manuscript, the authors used ChatGPT (OpenAI) for language editing and improvement of readability. The authors reviewed and revised all generated content and take full responsibility for the final manuscript.

## Supplementary Figure

**Supplementary Figure 1. Reactivity of psGFP in HEK293 cells.**

HEK293 cells expressing psGFP were stimulated with Na_2_S_4_ (0 or 200 μM) and imaged at indicated time points. Scale bar, 20 μm

**Supplementary Figure 2. Verification of SuTR in HEK293 cells.**

(A) HEK293 cells expressing SuTR were stimulated with Na_2_S_3_ (200 μM) and imaged at indicated time points. Scale bar, 20 μm (B) HEK293 cells were stimulated with Na_2_S_3_ (200 μM), or Na_2_S (200 μM). Data are represented as the mean ± SEM.

**Supplementary Figure 3. Leptomycin B does not inhibit the supersulfur-induced translocation of SuTR.**

(A) HeLa cells expressing SuTR were pre-treated with Leptomycin B (10 ng/ml) for 30 min, followed by treatment with Na_2_S_3_ (100 μM). Scale bar, 20 μm. (B) Quantification of the nuclear and cytoplasmic fluorescence intensity ratio of SuTR. Data are shown as the mean ± SEM from 15 cells. Significance was determined using two-way ANOVA with Sidak’s comparison test.

**Supplementary table 1.**
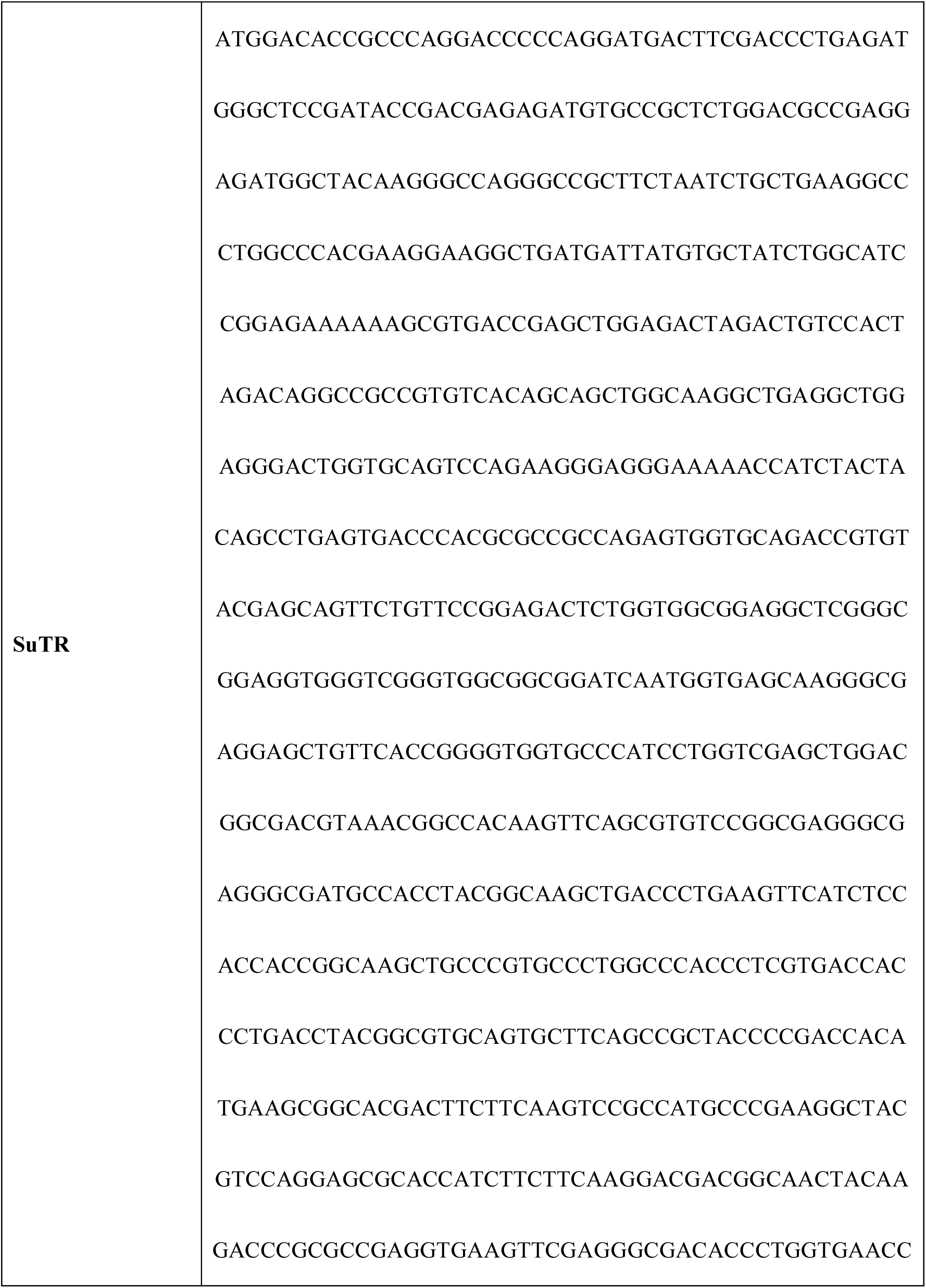
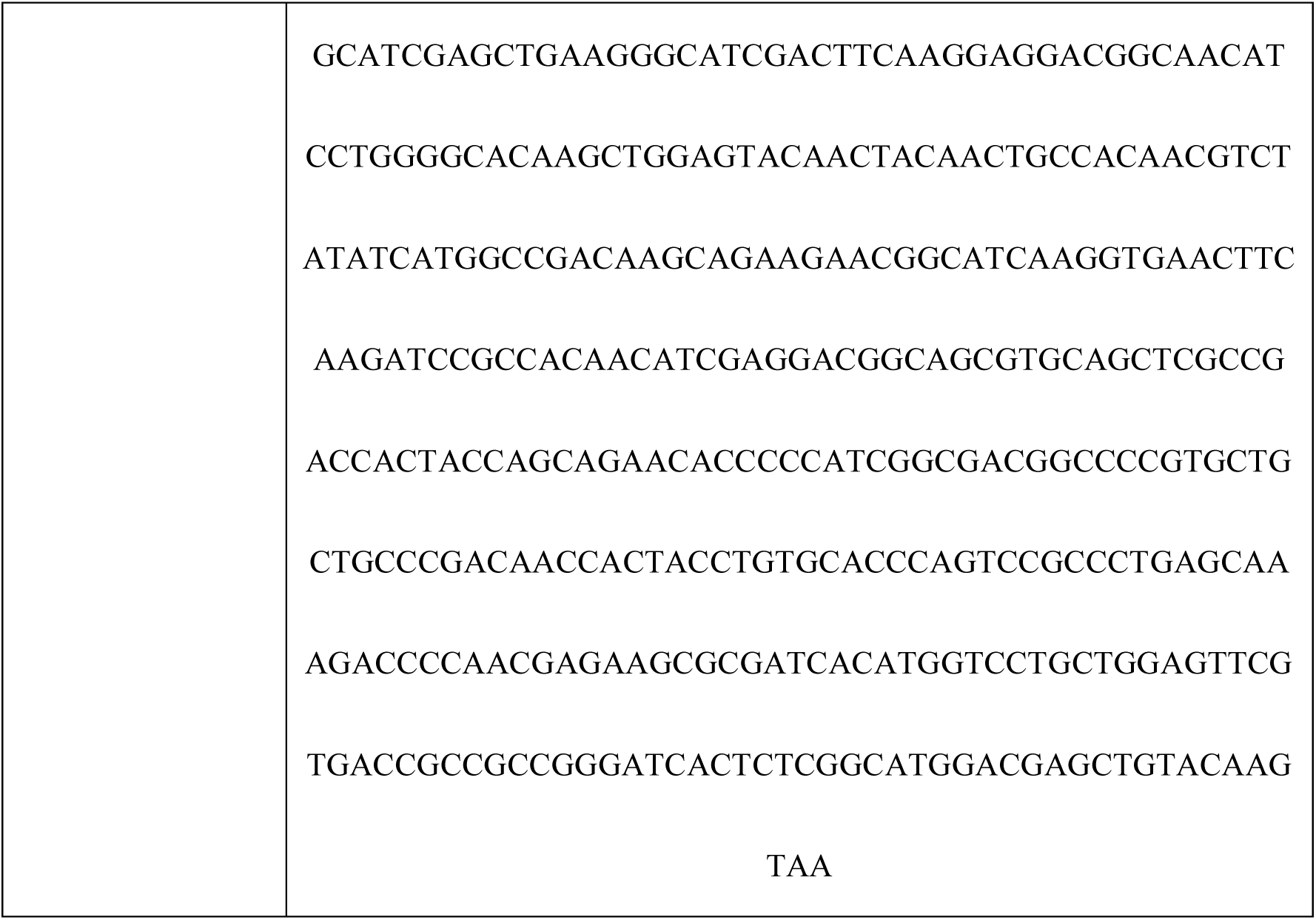
Nucleotide sequence of SuTR.

**Supplementary table 2.**
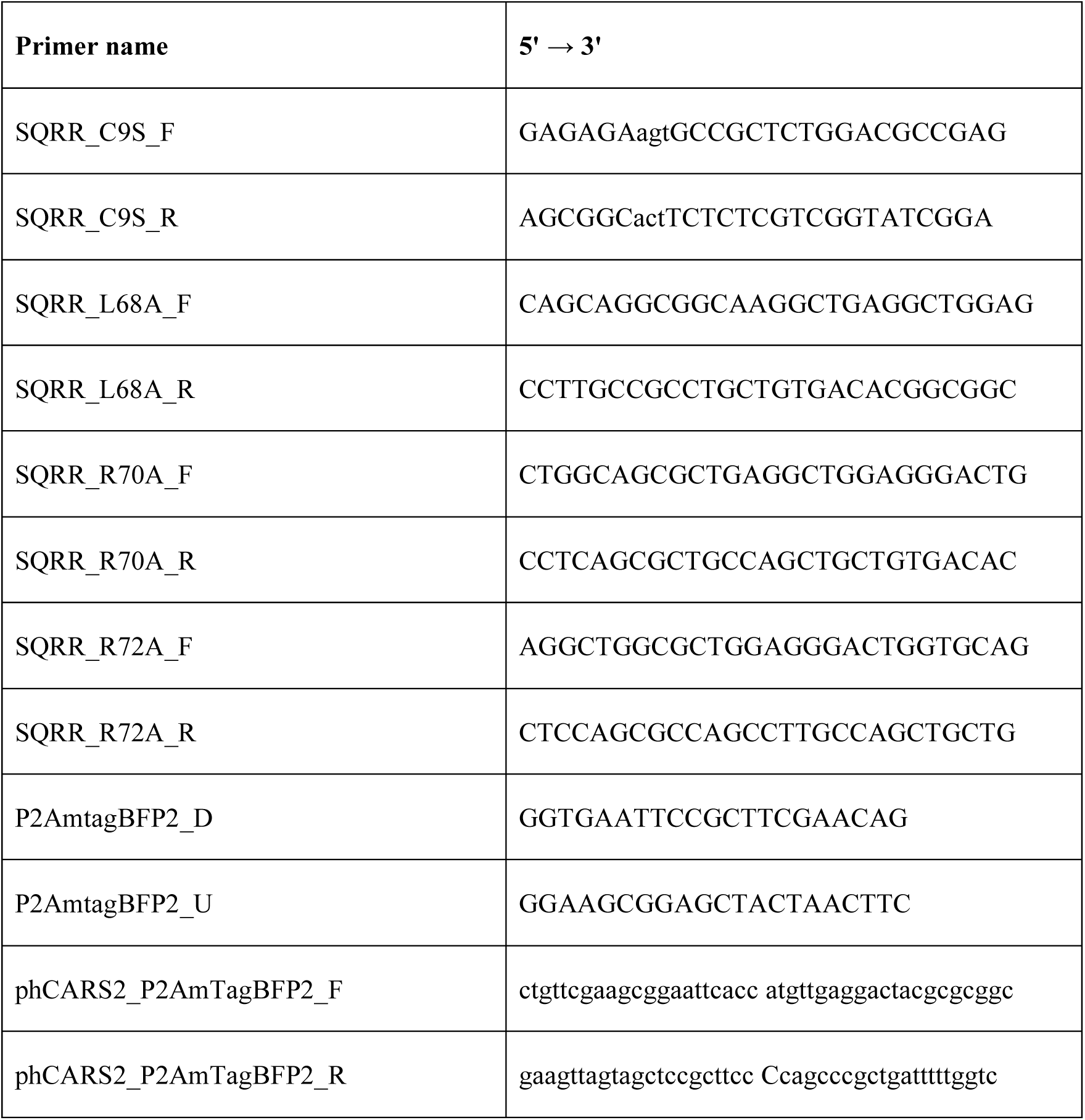
Primers.

## References

[1] U. Barayeu, T. Sawa, M. Nishida, F.Y. Wei, H. Motohashi, T. Akaike, Supersulfide biology and translational medicine for disease control, Br J Pharmacol 183 (1) (2026) 115–130. 10.1111/bph.16271.

[2] L. Zhou, A. Nishimura, K. Umezawa, Y. Kato, X. Mi, T. Ito, Y. Urano, T. Akaike, M. Nishida, Supersulfide catabolism participates in maladaptive remodeling of cardiac cells, J Pharmacol Sci 155 (4) (2024) 121–130. 10.1016/j.jphs.2024.05.002.

[3] H. Kimura, Signaling molecules: Hydrogen sulfide and polysulfide, Antioxidants and Redox Signaling 22 (5) (2015) 362–376. 10.1089/ars.2014.5869.

[4] H. Li, J. Li, C. Lü, Y. Xia, Y. Xin, H. Liu, L. Xun, H. Liu, FisR activates σ54-dependent transcription of sulfide-oxidizing genes in Cupriavidus pinatubonensis JMP134, Molecular Microbiology 105 (3) (2017) 373–384. 10.1111/mmi.13725.

[5] T.V. Mishanina, M. Libiad, R. Banerjee, Biogenesis of reactive sulfur species for signaling by hydrogen sulfide oxidation pathways, Nature Chemical Biology 11 (7) (2015) 457–464. 10.1038/nchembio.1834.

[6] M. Nishida, Y. Kumagai, H. Ihara, S. Fujii, H. Motohashi, T. Akaike, Redox signaling regulated by electrophiles and reactive sulfur species, Journal of Clinical Biochemistry and Nutrition 58 (2) (2016) 91–98. 10.3164/jcbn.15-111.

[7] T. Sawa, K. Ono, H. Tsutsuki, T. Zhang, T. Ida, M. Nishida, T. Akaike, Reactive Cysteine Persulphides: Occurrence, Biosynthesis, Antioxidant Activity, Methodologies, and Bacterial Persulphide Signalling, Advances in Microbial Physiology, 2018, pp. 1–28.

[8] A. Nishimura, K. Shimoda, T. Tanaka, T. Toyama, K. Nishiyama, Y. Shinkai, T. Numaga-Tomita, D. Yamazaki, Y. Kanda, T. Akaike, Y. Kumagai, M. Nishida, Depolysulfidation of Drp1 induced by low-dose methylmercury exposure increases cardiac vulnerability to hemodynamic overload, Sci Signal 12 (587) (2019). 10.1126/scisignal.aaw1920.

[9] A. Nishimura, S. Ogata, X. Tang, K. Hengphasatporn, K. Umezawa, M. Sanbo, M. Hirabayashi, Y. Kato, Y. Ibuki, Y. Kumagai, K. Kobayashi, Y. Kanda, Y. Urano, Y. Shigeta, T. Akaike, M. Nishida, Polysulfur-based bulking of dynamin-related protein 1 prevents ischemic sulfide catabolism and heart failure in mice, Nature Communications 16 (1) (2025) 276. 10.1038/s41467-024-55661-5.

[10] A. Nishimura, T. Tanaka, K. Shimoda, T. Ida, S. Sasaki, K. Umezawa, H. Imamura, Y. Urano, F. Ichinose, T. Kaneko, T. Akaike, M. Nishida, Non-thermal atmospheric pressure plasma-irradiated cysteine protects cardiac ischemia/reperfusion injury by preserving supersulfides, Redox Biology 79 (2025) 103445. 10.1016/j.redox.2024.103445.

[11] T. Akaike, T. Ida, F.Y. Wei, M. Nishida, Y. Kumagai, M.M. Alam, H. Ihara, T. Sawa, T. Matsunaga, S. Kasamatsu, A. Nishimura, M. Morita, K. Tomizawa, A. Nishimura, S. Watanabe, K. Inaba, H. Shima, N. Tanuma, M. Jung, S. Fujii, Y. Watanabe, M. Ohmuraya, P. Nagy, M. Feelisch, J.M. Fukuto, H. Motohashi, Cysteinyl-tRNA synthetase governs cysteine polysulfidation and mitochondrial bioenergetics, Nature Communications 8 (1) (2017). 10.1038/s41467-017-01311-y.

[12] C.M. Park, L. Weerasinghe, J.J. Day, J.M. Fukuto, M. Xian, Persulfides: current knowledge and challenges in chemistry and chemical biology, Molecular BioSystems 11 (7) (2015) 1775–1785. 10.1039/c5mb00216h.

[13] T. Zhang, T. Toyomoto, T. Sawa, T. Akaike, T. Matsunaga, Supersulfides: A Promising Therapeutic Approach for Autoinflammatory Diseases, Microbiol Immunol 69 (4) (2025) 191–202. 10.1111/1348-0421.13205.

[14] Y. Pan, T. Matsunaga, T. Zhang, T. Akaike, The Therapeutic Potential of Supersulfides in Oxidative Stress-Related Diseases, Biomolecules 15 (2) (2025) 172.

[15] É. Dóka, I. Pader, A. Bíró, K. Johansson, Q. Cheng, K. Ballagó, J.R. Prigge, D. Pastor-Flores, T.P. Dick, E.E. Schmidt, E.S.J. Arnér, P. Nagy, A novel persulfide detection method reveals protein persulfide- and polysulfide-reducing functions of thioredoxin and glutathione systems, Science Advances 2 (1) (2016). 10.1126/sciadv.1500968.

[16] T. Ida, T. Sawa, H. Ihara, Y. Tsuchiya, Y. Watanabe, Y. Kumagai, M. Suematsu, H. Motohashi, S. Fujii, T. Matsunaga, M. Yamamoto, K. Ono, N.O. Devarie-Baez, M. Xian, J.M. Fukuto, T. Akaike, Reactive cysteine persulfides and S-polythiolation regulate oxidative stress and redox signaling, Proceedings of the National Academy of Sciences of the United States of America 111 (21) (2014) 7606–7611. 10.1073/pnas.1321232111.

[17] J. Pan, K.S. Carroll, Persulfide reactivity in the detection of protein S-sulfhydration, ACS Chemical Biology 8 (6) (2013) 1110–1116. 10.1021/cb4001052.

[18] C.M. Park, I. Macinkovic, M.R. Filipovic, M. Xian, Use of the "Tag-Switch" Method for the Detection of Protein S-Sulfhydration, Methods in Enzymology, 2015, pp. 39–56.

[19] W. Chen, C. Liu, B. Peng, Y. Zhao, A. Pacheco, M. Xian, New fluorescent probes for sulfane sulfurs and the application in bioimaging, Chemical Science 4 (7) (2013) 2892–2896. 10.1039/c3sc50754h.

[20] K. Shimamoto, K. Hanaoka, Fluorescent probes for hydrogen sulfide (H2S) and sulfane sulfur and their applications to biological studies, Nitric Oxide - Biology and Chemistry 46 (2015) 72–79. 10.1016/j.niox.2014.11.008.

[21] K. Umezawa, M. Kamiya, Y. Urano, A Reversible Fluorescent Probe for Real-Time Live-Cell Imaging and Quantification of Endogenous Hydropolysulfides, Angewandte Chemie - International Edition 57 (30) (2018) 9346–9350. 10.1002/anie.201804309.

[22] X. Hu, H. Li, X. Zhang, Z. Chen, R. Zhao, N. Hou, J. Liu, L. Xun, H. Liu, Developing polysulfide-sensitive gfps for real-time analysis of polysulfides in live cells and subcellular organelles, Analytical Chemistry 91 (6) (2019) 3893–3901. 10.1021/acs.analchem.8b04634.

[23] Z. Li, Q. Wang, Y. Xia, L. Xun, H. Liu, A red fluorescent protein-based probe for detection of intracellular reactive sulfane sulfur, Antioxidants 9 (10) (2020) 1–13. 10.3390/antiox9100985.

[24] V. Haellman, T. Strittmatter, A. Bertschi, P. Stücheli, M. Fussenegger, A versatile plasmid architecture for mammalian synthetic biology (VAMSyB), Metab Eng 66 (2021) 41–50. 10.1016/j.ymben.2021.04.003.

[25] T. Tanida, K.I. Matsuda, S. Yamada, T. Hashimoto, M. Kawata, Estrogen-related Receptor β Reduces the Subnuclear Mobility of Estrogen Receptor α and Suppresses Estrogen-dependent Cellular Function*, Journal of Biological Chemistry 290 (19) (2015) 12332–12345. 10.1074/jbc.M114.619098.

[26] T. Shimizu, J. Shen, M. Fang, Y. Zhang, K. Hori, J.C. Trinidad, C.E. Bauer, D.P. Giedroc, S. Masuda, Sulfide-responsive transcriptional repressor SqrR functions as a master regulator of sulfide-dependent photosynthesis, Proceedings of the National Academy of Sciences 114 (9) (2017) 2355–2360. 10.1073/pnas.1614133114.

[27] J. Abramson, J. Adler, J. Dunger, R. Evans, T. Green, A. Pritzel, O. Ronneberger, L. Willmore, A.J. Ballard, J. Bambrick, S.W. Bodenstein, D.A. Evans, C.-C. Hung, M. O’Neill, D. Reiman, K. Tunyasuvunakool, Z. Wu, A. Þemgulytë, E. Arvaniti, C. Beattie, O. Bertolli, A. Bridgland, A. Cherepanov, M. Congreve, A.I. Cowen-Rivers, A. Cowie, M. Figurnov, F.B. Fuchs, H. Gladman, R. Jain, Y.A. Khan, C.M.R. Low, K. Perlin, A. Potapenko, P. Savy, S. Singh, A. Stecula, A. Thillaisundaram, C. Tong, S. Yakneen, E.D. Zhong, M. Zielinski, A. Žídek, V. Bapst, P. Kohli, M. Jaderberg, D. Hassabis, J.M. Jumper, Accurate structure prediction of biomolecular interactions with AlphaFold 3, Nature 630 (8016) (2024) 493-500. 10.1038/s41586-024-07487-w.

[28] N. Percie du Sert, V. Hurst, A. Ahluwalia, S. Alam, M.T. Avey, M. Baker, W.J. Browne, A. Clark, I.C. Cuthill, U. Dirnagl, M. Emerson, P. Garner, S.T. Holgate, D.W. Howells, N.A. Karp, S.E. Lazic, K. Lidster, C.J. MacCallum, M. Macleod, E.J. Pearl, O.H. Petersen, F. Rawle, P. Reynolds, K. Rooney, E.S. Sena, S.D. Silberberg, T. Steckler, H. Würbel, The ARRIVE guidelines 2.0: Updated guidelines for reporting animal research, British Journal of Pharmacology 177 (16) (2020) 3617–3624. 10.1111/bph.15193.

[29] M. Negrini, G. Wang, A. Heuer, T. Björklund, M. Davidsson, AAV Production Everywhere: A Simple, Fast, and Reliable Protocol for In-house AAV Vector Production Based on Chloroform Extraction, Curr Protoc Neurosci 93 (1) (2020) e103. 10.1002/cpns.103.

[30] S. Kasamatsu, T. Owaki, S. Komae, A. Kinno, T. Ida, T. Akaike, H. Ihara, Untargeted polysulfide omics analysis of alternations in polysulfide production during the germination of broccoli sprouts, Redox Biology 67 (2023) 102875. 10.1016/j.redox.2023.102875.

[31] M. Gutscher, A.-L. Pauleau, L. Marty, T. Brach, G.H. Wabnitz, Y. Samstag, A.J. Meyer, T.P. Dick, Real-time imaging of the intracellular glutathione redox potential, Nature Methods 5 (6) (2008) 553–559. 10.1038/nmeth.1212.

[32] M. Gutscher, M.C. Sobotta, G.H. Wabnitz, S. Ballikaya, A.J. Meyer, Y. Samstag, T.P. Dick, Proximity-based Protein Thiol Oxidation by H2O2-scavenging Peroxidases*♦, Journal of Biological Chemistry 284 (46) (2009) 31532–31540. 10.1074/jbc.M109.059246.

[33] T. Akaike, T. Ida, F.-Y. Wei, M. Nishida, Y. Kumagai, M.M. Alam, H. Ihara, T. Sawa, T. Matsunaga, S. Kasamatsu, A. Nishimura, M. Morita, K. Tomizawa, A. Nishimura, S. Watanabe, K. Inaba, H. Shima, N. Tanuma, M. Jung, S. Fujii, Y. Watanabe, M. Ohmuraya, P. Nagy, M. Feelisch, J.M. Fukuto, H. Motohashi, Cysteinyl-tRNA synthetase governs cysteine polysulfidation and mitochondrial bioenergetics, Nature Communications 8 (1) (2017) 1177. 10.1038/s41467-017-01311-y.

[34] D.A. Capdevila, B.J.C. Walsh, Y. Zhang, C. Dietrich, G. Gonzalez-Gutierrez, D.P. Giedroc, Structural basis for persulfide-sensing specificity in a transcriptional regulator, Nat Chem Biol 17 (1) (2021) 65–70. 10.1038/s41589-020-00671-9.

[35] G.T. Hanson, R. Aggeler, D. Oglesbee, M. Cannon, R.A. Capaldi, R.Y. Tsien, S.J. Remington, Investigating Mitochondrial Redox Potential with Redox-sensitive Green Fluorescent Protein Indicators*, Journal of Biological Chemistry 279 (13) (2004) 13044–13053. 10.1074/jbc.M312846200.

[36] D.M. Veine, L.D. Arscott, C.H. Williams, Redox Potentials for Yeast, Escherichia coli and Human Glutathione Reductase Relative to the NAD+/NADH Redox Couple: Enzyme Forms Active in Catalysis, Biochemistry 37 (44) (1998) 15575–15582. 10.1021/bi9811314.

[37] T. Gosselin-Monplaisir, B. Enjalbert, S. Uttenweiler-Joseph, J.C. Portais, S. Heux, P. Millard, Overflow metabolism in bacterial, yeast, and mammalian cells: different names, same game, Mol Syst Biol 21 (11) (2025) 1419–1433. 10.1038/s44320-025-00145-x.

[38] S. Regot, J.J. Hughey, B.T. Bajar, S. Carrasco, M.W. Covert, High-sensitivity measurements of multiple kinase activities in live single cells, Cell 157 (7) (2014) 1724–1734. 10.1016/j.cell.2014.04.039.

[39] T. Kudo, S. Jeknić, D.N. Macklin, S. Akhter, J.J. Hughey, S. Regot, M.W. Covert, Live-cell measurements of kinase activity in single cells using translocation reporters, Nature Protocols 13 (1) (2018) 155–169. 10.1038/nprot.2017.128.

[40] S.J. Tsai, Y. Gong, A. Dabbs, F. Zahra, J. Xu, A. Geske, M.J. Caterina, S.J. Gould, Enhanced kinase translocation reporters for simultaneous real-time measurement of PKA, ERK, and calcium, J Biol Chem 301 (3) (2025) 108183. 10.1016/j.jbc.2025.108183.

[41] B. Saykali, A.D. Tran, J.A. Cornwell, M.A. Caldwell, P.R. Sangsari, N.Y. Morgan, M.J. Kruhlak, S.D. Cappell, S. Ruiz, Lineage-specific CDK activity dynamics characterize early mammalian development, Cell Rep 44 (4) (2025) 115558. 10.1016/j.celrep.2025.115558.

[42] D.V. Chandrashekar, B.N. DuBois, M. Rashid, R. Mehvar, Effects of chronic cirrhosis induced by intraperitoneal thioacetamide injection on the protein content and Michaelis-Menten kinetics of cytochrome P450 enzymes in the rat liver microsomes, Basic Clin Pharmacol Toxicol 132 (2) (2023) 197–210. 10.1111/bcpt.13813.

[43] A.A. Mohamed, A.A. Shoun, R.A. El-Kadi, S.O. Abd El-Maseh, S.A. Abass, Herbal management of TAA-induced liver toxicity: Fibrosis, cirrhosis, and hepatocellular carcinoma, Pathology - Research and Practice 273 (2025) 156141. 10.1016/j.prp.2025.156141.

[44] K. Honda, T. Hishiki, S. Yamamoto, T. Yamamoto, N. Miura, A. Kubo, M. Itoh, W.-Y. Chen, M. Takano, T. Yoshikawa, T. Kasamatsu, S. Sonoda, H. Yoshizawa, S. Nakamura, Y. Itai, M. Shiota, D. Koike, M. Naya, N. Hayakawa, Y. Naito, T. Matsuura, K. Iwaisako, T. Masui, S. Uemoto, K. Nagashima, Y. Hashimoto, T. Sakuma, O. Matsubara, W. Huang, T. Ida, T. Akaike, Y. Masugi, M. Sakamoto, T. Kato, Y. Ino, H. Yoshida, H. Tsuda, N. Hiraoka, Y. Kabe, M. Suematsu, On-tissue polysulfide visualization by surface-enhanced Raman spectroscopy benefits patients with ovarian cancer to predict post-operative chemosensitivity, Redox Biology 41 (2021) 101926. 10.1016/j.redox.2021.101926.

[45] M. Shiota, M. Naya, T. Yamamoto, T. Hishiki, T. Tani, H. Takahashi, A. Kubo, D. Koike, M. Itoh, M. Ohmura, Y. Kabe, Y. Sugiura, N. Hiraoka, T. Morikawa, K. Takubo, K. Suina, H. Nagashima, O. Sampetrean, O. Nagano, H. Saya, S. Yamazoe, H. Watanabe, M. Suematsu, Gold-nanofève surface-enhanced Raman spectroscopy visualizes hypotaurine as a robust anti-oxidant consumed in cancer survival, Nature Communications 9 (1) (2018). 10.1038/s41467-018-03899-1.

