## Supplementary figures and images for "Development of a genetically encoded supersulfide-dependent translocation reporter"

### Supplemental figure 1

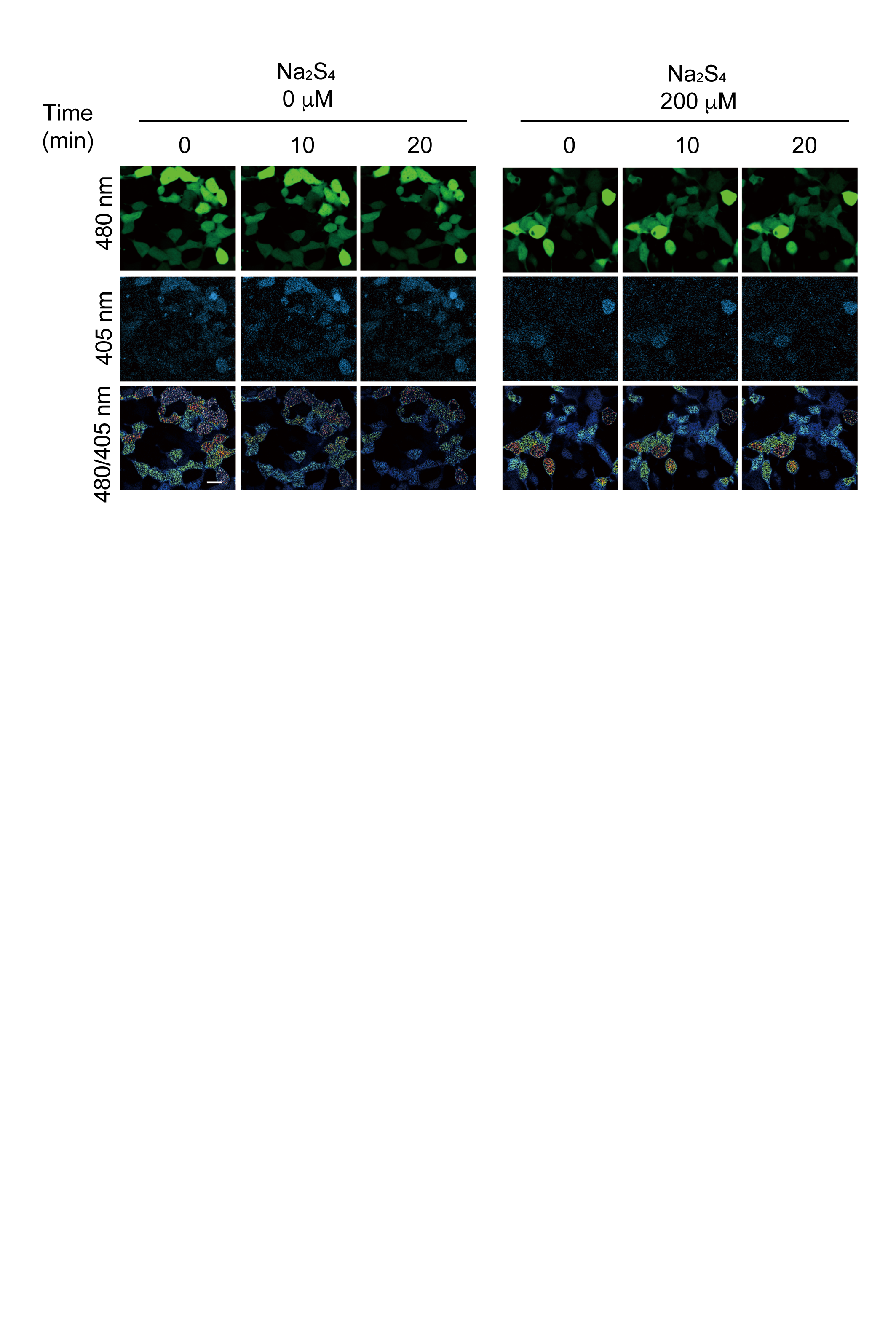

### Supplemental figure 2

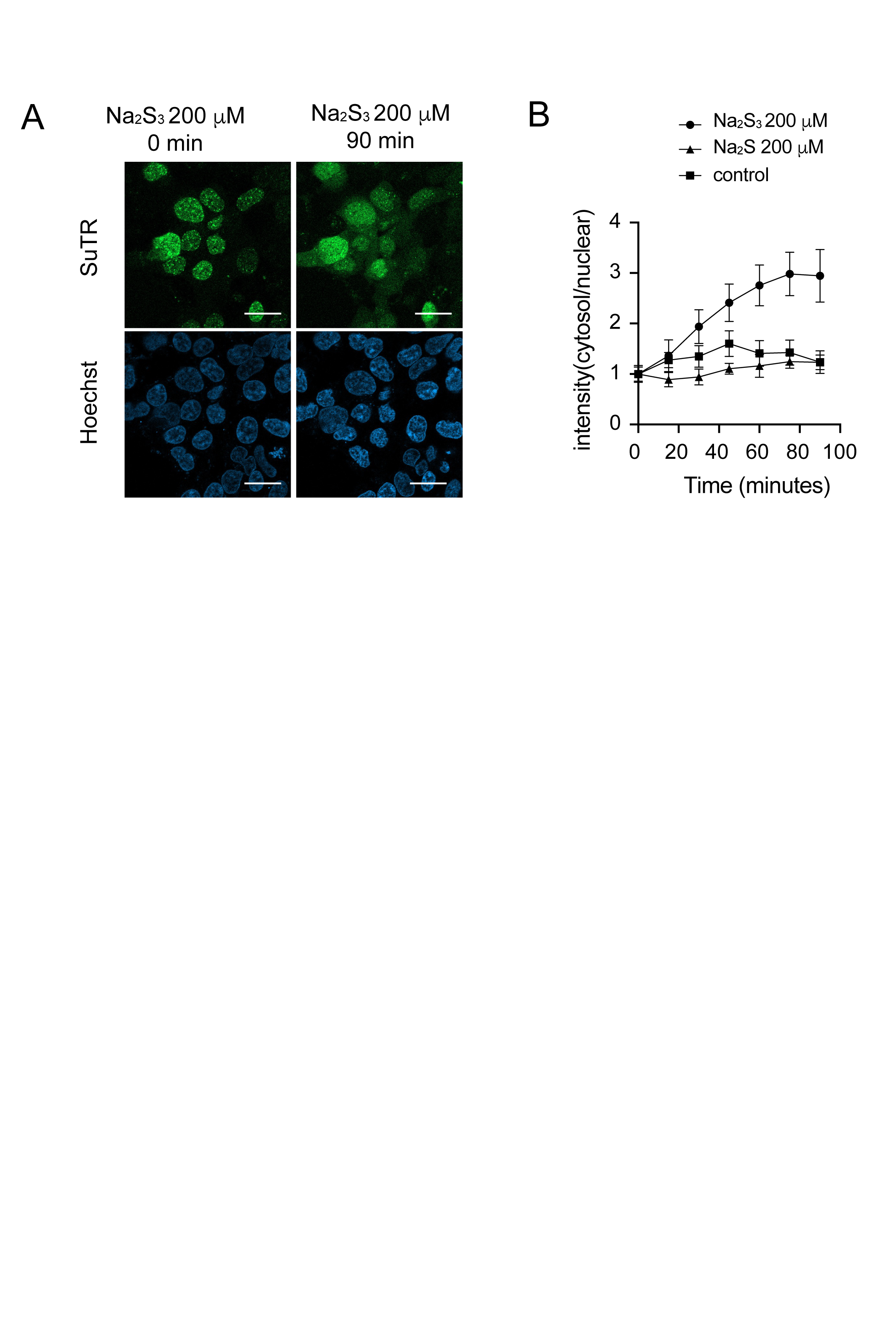

### Supplemental figure 3

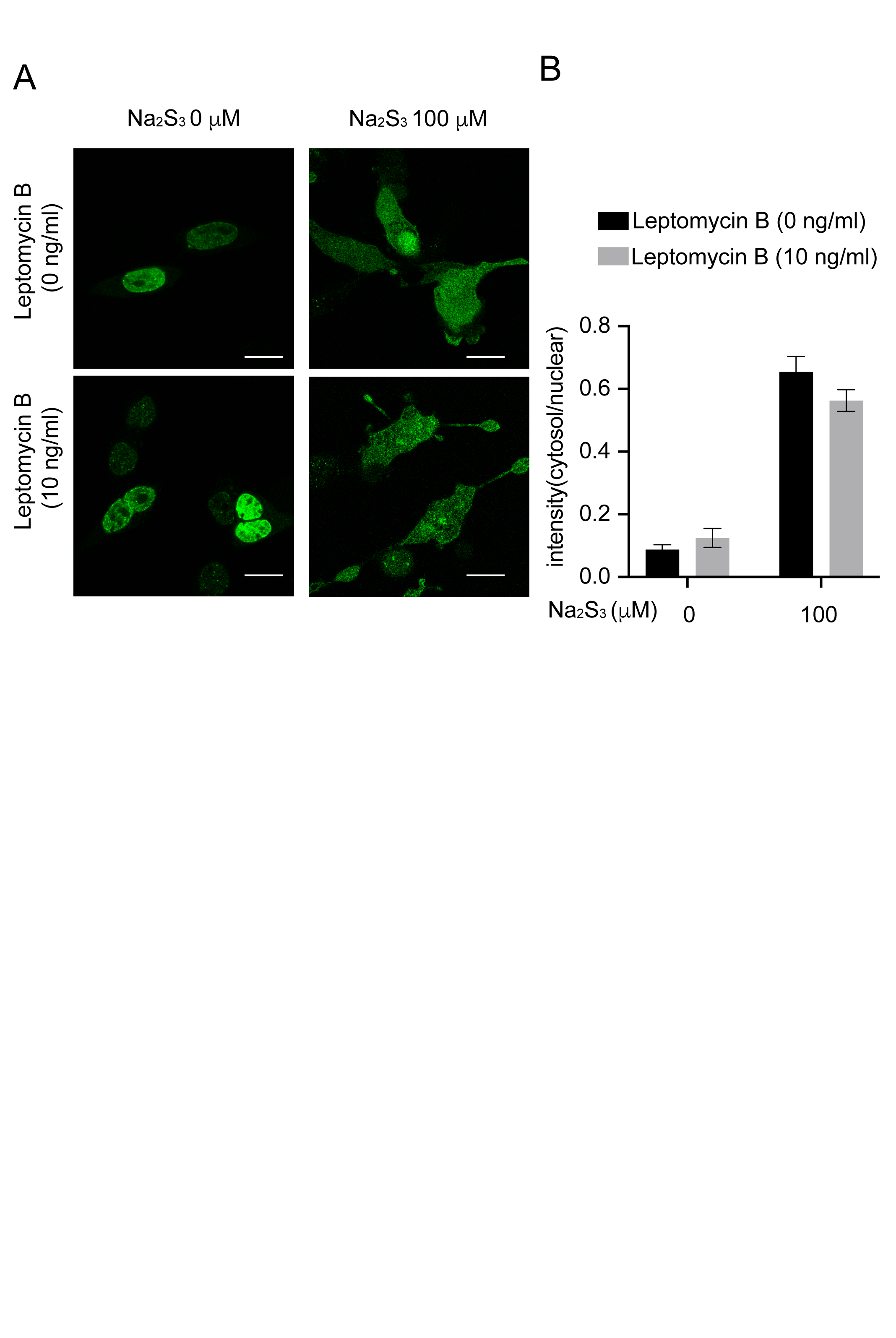
